# Cannabinoids differentially modulate behavioral and developmental responses to ethanol in *Drosophila*

**DOI:** 10.1101/2021.08.14.456340

**Authors:** Jianzheng He, Si Yun Ng, Alice Mei Xien Tan, Wei Lin Yong, Fengwei Yu

**Author notes:** These authors contributed equally to this work (Jianzheng He, Si Yun Ng).

## Abstract

Prolonged prenatal or adult exposure to ethanol is detrimental to mental and physical well-being, resulting in developmental abnormalities, progressive addiction and ultimate death. A growing number of studies have shown the therapeutic potential of cannabinoids in ethanol-related behaviors in mammals. However, the potential pharmacological actions of cannabinoids in ethanol responses have not been examined in the model organism *Drosophila melanogaster*. Here, we systematically investigated the effects of various cannabinoids on ethanol preference, ethanol sensitivity and tolerance, and ethanol-induced developmental defect in *Drosophila*. We showed that treatment with the phytocannabinoid cannabidiol (CBD) displayed a significant decrease in preference for consuming ethanol in adult flies. Interestingly, cannabinoids exhibited differential roles in short- and long-term ethanol tolerance in flies. Although cannabinoids had no detectable effects on short-term ethanol tolerance, CBD and the endocannabinoid anandamide (AEA) suppressed long-term tolerance to ethanol. Moreover, ethanol exposure delayed larval-to-pupal development and increased larval/pupal size. Unexpectedly, treatment with CBD or endocannabinoids did not attenuate ethanol-induced developmental delay, instead, exacerbated its detrimental effect. Thus, our systematical study reveals, for the first time, a differential role of the cannabinoids in the modulation of ethanol-related responses in *Drosophila*.

## Introduction

*Cannabis sativa*, also known as marijuana or hemp, is an annual herbaceous plant known for its medicinal and recreational purposes. Since the discovery of the main constituents in *Cannabis* in the 1960s and the naturally-occurring endocannabinoid system in 1990s (Mechoulam et al., 2014), researchers have focused on understanding of the medicinal relevance of different constituents derived from *Cannabis* and endogenous endocannabinoids. The *Cannabis sativa* plant produces more than 545 constituents (ElSohly and Gul, 2015), among which more than 100 constituents were isolated and identified as phytocannabinoids (Hanuš et al., 2016). Aside from the known psychoactive component Δ^9^-tetrahydrocannabinol (THC), cannabidol (CBD) is the most commonly studied phytocannabinioid due to its non-psychotomimetic property. Other non-psychoactive phytocannabinoids including cannabigerol (CBG), cannabichromene (CBC) and cannabidivarin (CBDV) have also been characterized. The plant derivatives or synthetic cannabinoids interact with the endocannabinoid system which consists of endogenous ligands N-arachidonoylethanolamine (AEA) and 2-arachidonoylglycerol (2-AG), cannabinoid receptors and endocannabinoid-degrading enzymes (Mechoulam et al., 2014). Collectively, these cannabinoids are particularly of high interest due to their potential therapeutic values. Preclinical studies in rodents showed that CBD decreased alcohol intake with improvement in hepatic and neurocognitive outcomes (De Ternay et al., 2019; Turna et al., 2019). Recently, a clinical study reported reduction in disruptive behaviors in children and young adults treated with CBD (Koren et al., 2021). Furthermore, emerging studies in *Drosophila* have documented the therapeutic effects of cannabinoids in food intake and various diseases including cardiovascular disease, Parkinson’s disease and seizure. Cannabinoids exhibited an inhibitory effect on food intake presumably through reduced lipid metabolism (He et al., 2021). Prolonged inhalation of marijuana increased cardiac contractility in adult flies (Gómez et al., 2019). The neuroprotective role of synthetic cannabinoid CP55940 and endocannabinoids was evident in paraquat-induced toxicity (Jimenez-Del-Rio et al., 2008) and seizure (Jacobs and Sehgal, 2020), respectively. Although a range of cannabinoids have been found to modulate diverse behavioral effects in *Drosophila*, the canonical CB_1/2_ receptors were found to be non-existent (McPartland et al., 2001). It is plausible that cannabinoids exert their effects through non-canonical G-protein coupled receptors (GPCR) which consist of 5HT_1A_ receptors, GPR55 and transient receptor potential (TRP) channels (Cristino et al., 2020). These studies are suggestive of cannabinoids acting on these non-canonical targets to modulate various behaviors in these pathological states.

Fetal alcohol spectrum disorder (FASD) and alcohol use disorder (AUD) are major global health issues resulting from increased alcohol consumption driven by social-cultural factors (Carvalho et al., 2019; Lange et al., 2017). While human studies and rodent models are popular among researchers to investigate alcohol-related disorders (Ehrhart et al., 2019; Marquardt and Brigman, 2016), *Drosophila melanogaster* has also been increasingly used as an important model to elucidate the underlying mechanisms due to the high conservation of genome and resemblance of alcohol-induced behaviors to mammals. It has been well documented that the adult flies exhibited the key features of alcohol-induced behaviors. The fruit flies were initially found to be attracted to ethanol with the ability to detect the presence of ethanol (McKenzie and McKechnie, 1979) and more recent studies revealed that flies displayed preference to ethanol food, a behavioral response similar to humans (Devineni and Heberlein, 2009; Ja et al., 2007; Park and Ja, 2020). Acute and chronic ethanol intoxication led to an increase in locomotion in the flies (Wolf et al., 2002) which progressed to impairment of motor skills with the loss of postural control and the inability to upright themselves (Moore et al., 1998). This was followed by subsequent sedation (Petruccelli et al., 2016; Singh and Heberlein, 2000; Wolf et al., 2002). When flies were repeatedly exposed to ethanol upon recovery, the flies developed tolerance to ethanol with increased latency to sedation (Parkhurst et al., 2018; Scholz et al., 2005; Scholz et al., 2000). Prolonged exposure to ethanol or ethanol at high concentrations could result in lethality (Devineni and Heberlein, 2012). To mimic prenatal exposure to ethanol in FASD, involuntary feeding of the larvae with ethanol exhibited developmental delay to eclosion of adult flies which displayed altered neurobehavioral changes (McClure et al., 2011).

Here, we characterized the role of cannabinoids in the modulation of various ethanol responses using the *Drosophila melanogaster* model. We confirmed that the adult flies exhibited preferential ethanol consumption, ethanol tolerance, and ethanol-induced developmental delay, consistent with the previous studies. Interestingly, treatment with CBD significantly decreased ethanol preference in adult flies. Moreover, we showed that flies treated with CBD and AEA displayed less resistance to ethanol sedation with suppression of long-term tolerance to ethanol. In addition, we reported unexpected findings that ethanol-induced developmental delay was not attenuated by CBD or endocannabinoids treatment. Overall, our study systematically interrogates the effects of cannabinoids on various ethanol-related behaviors and suggests selectivity of cannabinoids on ethanol responses.

## Material and methods

### Fly stocks and maintenance

The *Canton S* flies (#64349) used in all experiments were obtained from the Bloomington *Drosophila* stock center (BDSC), Indiana, USA. All flies were raised on standard food medium under a 12 h light/12 h dark cycle with temperature maintained at 25°C and relative humidity at 70%. All behavioral assays were performed on 3-5-day-old adult male flies and these flies were subjected to light CO_2_ anesthesia and allowed to recover for at least 2 days before any behavioral experimentation.

### Drug treatment and delivery

The following cannabinoids were used in this study (Table 1): 1) phytocannabinoids – CBD (0.02 mg/ml, 0.1 mg/ml, 0.5 mg/ml and 1 mg/ml), CBDV, CBG and CBC (0.1 mg/ml), 2) endocannabinoids – 2-AG and AEA (0.01 mg/ml and 0.1 mg/ml), and 3) synthetic cannabinoids - CP55940 (0.02 mM, 0.05 mM and 0.5 mM). These drugs were dissolved in ethanol or methanol and the stock solutions were further diluted to the desired concentrations in different food medium depending on behavioral assays. Due to the nature of behavioral tests used in this study, various feeding methods were utilized to treat the flies.

**Table 1.** The list of cannabinoids used in this study.

| **Cannabinoids** | **Name** | **Solvent** | **Final Concentration** | **Source** |
| --- | --- | --- | --- | --- |
| Phytocannabinoids | Cannabidiol (CBD) | Methanol | 0.02 mg/ml<br>0.1 mg/ml<br>0.5 mg/ml<br>1 mg/ml | Cayman Chemicals, USA |
|  | Cannabidivarin (CBDV) | Methanol | 0.1 mg/ml |  |
|  | Cannabichromene (CBC) | Methanol | 0.1 mg/ml |  |
|  | Cannabigerol (CBG) | Methanol | 0.1 mg/ml |  |
| Endocannabinoids | Anandamide (AEA) | Ethanol | 0.01 mg/ml<br>0.1 mg/ml | Tocris Bioscience, UK |
|  | 2-Arachidonoylglycerol (2-AG) | Methanol | 0.01 mg/ml<br>0.1 mg/ml |  |
| Synthetic cannabinoids | CP55940 (mimics the effects of $\Delta^9$-THC) | Methanol | 0.02 mM (0.008 mg/ml)<br>0.05 mM (0.02 mg/ml)<br>0.5 mM (0.2 mg/ml) | Sigma-Aldrich, USA |

For the ethanol preference assay, flies were presented with cannabinoid-containing liquid food (5% sucrose, 5% yeast extract) or the respective control food in the glass capillaries while cannabinoid-containing solid food (2% agar and 10% sucrose) was administered in the caps of Eppendorf tubes to assess ethanol sedation sensitivity and tolerance, and ethanol-induced lethality in adult flies. In the ethanol developmental exposure assay, eggs were collected and placed on standard food medium containing cannabinoids and/or ethanol.

### Ethanol preference assay

The preference for consuming ethanol was assessed using the CApillary FEeder (CAFE) assay (Devineni and Heberlein, 2009; Ja et al., 2007). Eight adult male flies were transferred into a vial containing 1% agarose with four 5 μl glass capillaries (VWR, USA). The capillaries were inserted from the top of the vial via adaptors made of pipette tips (He et al., 2021). Following pre-treatment with cannabinoid-containing liquid food or control liquid food for two days, the flies were presented with two capillaries filled with normal liquid food and the remaining with normal liquid food + 15% ethanol. The amount of food consumed in the capillaries was measured for two consecutive days and replaced with fresh food daily. Identical vials lacking flies were utilized as controls in each assay to determine evaporation from the capillaries. The mean amount of evaporation was subtracted from the food consumption data. The ethanol preference index was quantified as the (consumption of food with ethanol – consumption of normal liquid food) / total consumption.

### Ethanol sedation sensitivity and tolerance assays

The ethanol sedation sensitivity and tolerance assays were performed as previously described (Maples and Rothenfluh, 2011). On the experimental day, the flies were transferred to new empty vials which were subsequently covered with a fresh plug infused with 500 µl of 75% ethanol. The vials were tapped three times on the lab bench at a 1-minute interval to startle and knock the flies to the bottom of the vial. Following a 10-second observational period, the number of sedated flies at the bottom of the vial that were unable to upright themselves was recorded every minute. The time to 50% sedation (ST50) was determined whereby 50% of the flies remained sedated. To examine the effect of cannabinoids on ethanol sedation sensitivity and short-term ethanol tolerance, groups of 8 male flies underwent cannabinoid training for 2 or 4 days before behavioral testing. The sensitivity to ethanol-induced sedation of the flies was measured as ST50 following the first exposure to ethanol vapor for 25 minutes (ST50-0hr). Flies were subsequently transferred to normal food vials to recover at 25°C. Flies were examined for ethanol sedation (ST50-4hr) four hours following the first ethanol exposure. The short-term ethanol tolerance of the flies was assessed through the percentage change in time taken for ethanol sedation at four hours (ST50-4hr – ST50-0hr) / ST50-0hr (%). To further investigate the effects of cannabinoids on long-term ethanol tolerance, the flies were subjected to repeated ethanol exposure for 25 minutes daily for 5 days. Following the assessment of the innate ethanol tolerance of flies on day 0 prior to cannabinoid treatment, the flies were treated with cannabinoids for 4 days and the behavioral assay was performed daily concomitantly with cannabinoid treatment. The ST50 was measured every 24 hours for 4 consecutive days to determine the long-term change in ethanol tolerance. Long-term tolerance to ethanol was calculated by determining the percentage change in ST50 at daily (ST50-Day1-4 – ST50-Day0) / ST50-Day0 (%), respectively.

### Ethanol lethality assay

To assess whether cannabinoids alter ethanol-induced lethality, the flies were placed into an empty vial with a fresh plug containing 1 ml of 100% ethanol for 45 minutes following pre-treatment with cannabinoids for 2 or 4 days. After exposure to the ethanol vapor, they were transferred into vials with normal food and allowed to recover for 24 hours. The number of dead flies was counted. The ethanol-induced lethality of the flies (%) was calculated as (number of dead flies/number of total flies) x 100%.

### Developmental ethanol exposure assay

Eggs were collected on standard fly food medium for 24 hours. These eggs were suspended in sterile water and the same volume of eggs was distributed to different vials containing food medium with 1) ethanol (0%, 5%, 7.5% and 10%), 2) cannabinoids (0.02 mg/ml, 0.1 mg/ml and 0.5 mg/ml CBD, 0.02 mM CP55940, 0.01 mg/ml AEA and 0.01 mg/ml 2-AG), 3) ethanol (10%) + cannabinoids. The number of pupae was counted every 12 h from 5 to 12 days AEL and these data was used to calculate the time taken for the development of 50% of the eggs to pupa formation (DT50) and the egg-to-pupa survival. To examine whether ethanol affects larval and pupa growth, the size of the wondering third instar larvae (wL3) and pupa was measured using ImageJ.

### Statistical Analysis

Statistical analysis was performed using Prism 7.03 (GraphPad Software, USA). Data was analyzed using Student’s t-test or one-way ANOVA followed by Dunnett’s post-hoc test. When one-way ANOVA indicates significance with Bartlett’s test which represents unequal variance, non-parametric Kruskal-Wallis test followed by Dunn’s multiple comparison test was performed. Differences between the treatment groups were considered to be statistically significant at * p < 0.05, ** p < 0.01, *** p < 0.001. All graphs were represented as mean ± standard error of mean (S.E.M).

## Results

### CBD decreased preference for ethanol in flies

A previous study demonstrated that adult flies exhibit preferential ethanol consumption (Devineni and Heberlein, 2009), indicating that *Drosophila melanogaster* can successfully model this specific behavior which is conserved in vertebrates. Consistently, our findings showed that the control flies exhibited a robust preference for consuming the food containing a high concentration of ethanol (15%) when compared to non-ethanol food (Figure 1). The flies were pre-trained with cannabinoids-containing liquid food in the capillaries for 2 days (day-2 and day-1) prior to the measurement of ethanol preference in the subsequent two days (day 1 and day 2) using the CAFE assay (Figure 1A). We investigated whether cannabinoids could affect ethanol preference in these pre-treated adult flies. Amongst the phytocannabinoids, pre-treatment with 0.1 mg/ml and 1mg/ml CBD significantly reduced the ethanol preference index on day 2, but not on day 1 (Figures 1B and 1C). In line with our previous study (He et al., 2021), the average food consumption of flies was significantly decreased with CBD at the higher concentration (Figures S1B), but not the low concentration (Figure S1A). In addition, we also assessed the preference for ethanol with various phytocannabinoids including CBDV, CBC and CBG at 0.1 mg/ml. In contrast to CBD, these flies displayed similar preference for ethanol on both day 1 and day 2 to their respective controls (Figures 1D-1F). Likewise, the ethanol preference of flies pre-treated with the synthetic cannabinoid CP55940 was not altered (Figures 1G) while a significant reduction in food intake was observed in CP55940-treated flies (Figure S1C). Pre-treatment with endocannabinoids AEA (Figure 1H) and 2-AG (Figure 1I) also did not affect ethanol preference in flies. However, a trend towards decreased preference of ethanol with 2-AG treatment was observed (Figure 1I). Taken together, these findings suggest an inhibitory role of CBD on ethanol preference in flies.

**Figure 1.**
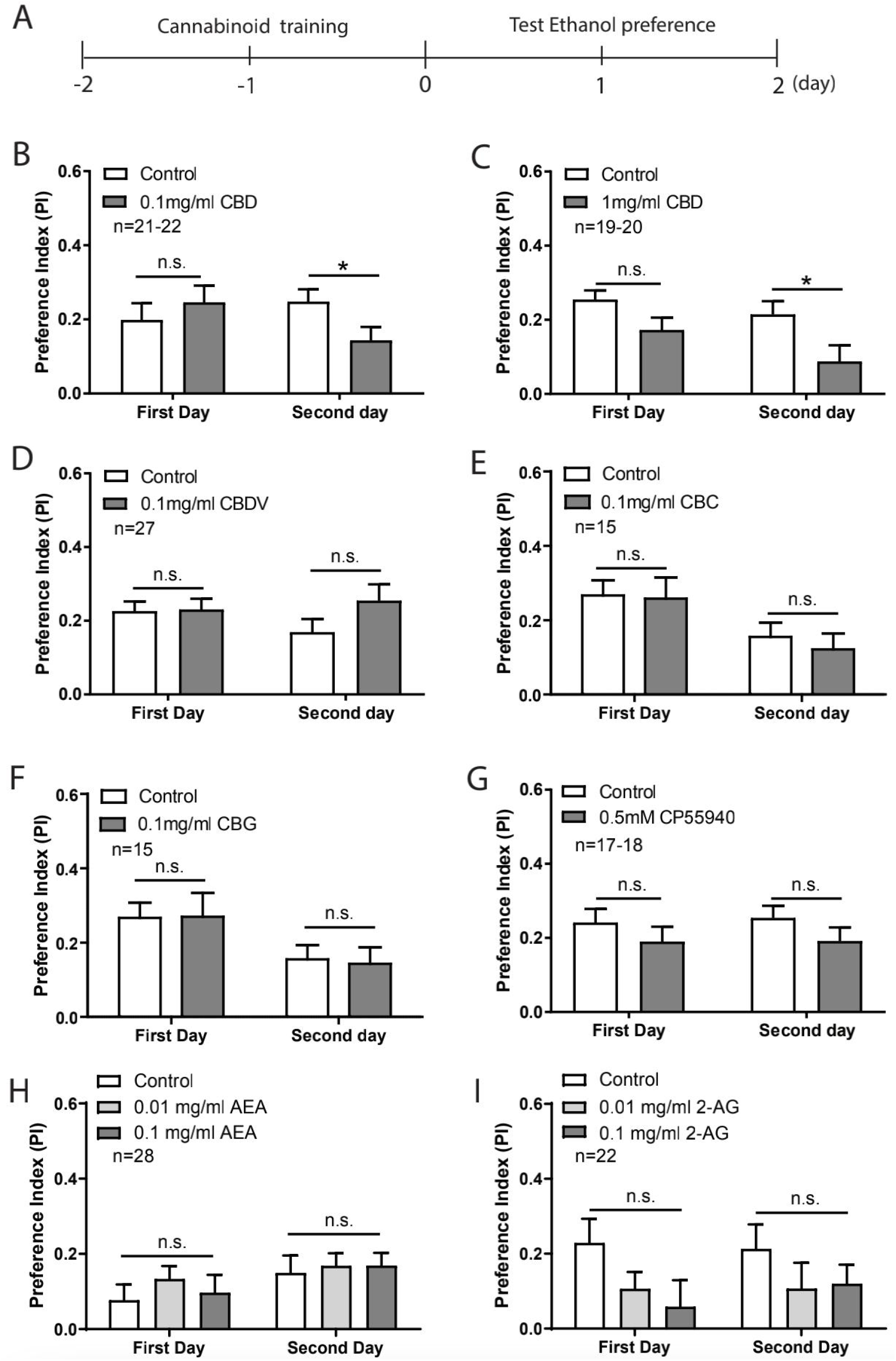
CBD decreased ethanol preference in flies. (A) A schematic diagram of the experimental timeline for the ethanol preference assays. Following two days of cannabinoid treatment, flies were fed with control or ethanol liquid food and the preference for ethanol was measured daily for two consecutive days (day 1 and day 2). (B, C) Pre-treatment of 0.1 mg/ml CBD (B) and 1 mg/ml CBD (C) significantly decreased preference for ethanol food on day 2. The ethanol preference was not affected in flies pre-treated with 0.1 mg/ml CBDV (D), 0.1 mg/ml CBC (E), 0.1 mg/ml CBG (F), 0.5 mM CP55940 (G) (n=15-27). The ethanol preference of flies pre-treated with AEA (H) was also similar to the control groups (n=28). Flies fed with 2-AG (I) displayed a trend of reduction in ethanol preference although statistically non-significant (n=22). Significance was determined using two-tailed unpaired t-test and one-way ANOVA, * p < 0.05. All data are represented as mean ± S.E.M.

### Cannabinoids did not alter short-term ethanol tolerance in flies

Similar to ethanol preference, the flies exhibit a series of acute ethanol intoxicated behaviors which are conserved in mammals. The acute reversible events begin with hyperactivity, followed by uncoordinated motor activity and subsequent sedation (Bainton et al., 2000; Singh and Heberlein, 2000). We first examined whether cannabinoids could regulate ethanol sedation sensitivity in flies. The flies were pre-treated with cannabinoid-containing food for 2 or 4 consecutive days prior to the measurement of sedation sensitivity to ethanol at two time points of 0 hr and 4 hr (Figure 2A and S2A). The ethanol-induced sedation sensitivity was quantified as the time taken for 50% of the flies to be sedated during ethanol exposure (ST50). Flies were pre-treated for 2 days with various phytocannabinoids including CBD at 0.1 mg/ml and 1 mg/ml (Figure 2B), CBC, CBDV (Figure 2D), CBG (Figure 2F) at 0.1 mg/ml, synthetic cannabinoid CP55940 at 0.5 mM (Figure 2F) and endocannabinoids AEA (Figure 2H) and 2-AG (Figure 2J) at 0.01 mg/ml and 0.1 mg/ml. The ST50s of the flies treated with these cannabinoids were similar as compared to the respective controls. Thus, these cannabinoids did not affect ethanol sedation sensitivity in flies. Prolonged 4-day treatment of selective cannabinoids consisting of CBD (Figure S2B), CP55940 (Figure S2D) and 2-AG (Figure S2F) did not alter the sensitivity to ethanol either. We further investigated whether cannabinoids could regulate short-term tolerance to ethanol which is derived from the percentage change in sedation sensitivity at two different time points. Previous studies provided evidence of the development of tolerance to ethanol in adult flies during the second ethanol exposure (Berger et al., 2008; Scholz et al., 2000). Consistently, we also found that flies pre-treated with the control solutions were more resistant to sedation during the second exposure to ethanol with an increase in tolerance change of 85.84 ± 4.71 %. CBD treatment at both concentrations led to a trend towards decreased short-term ethanol tolerance; however, it is not statistically significant (Figure 2C). Furthermore, flies pre-treated with CBD for 4 days did not alter the short-term ethanol tolerance (Figure S2C). Two-day treatment with the other phytocannabinoids including 0.1 mg/ml CBC, 0.1 mg/ml CBDV or 0.1 mg/ml CBG did not affect short-term ethanol tolerance (Figure 2E, 2G). Similarly, flies pre-treated with CP55940 at 0.5 mM or AEA at 0.01 mg/ml and 0.1 mg/ml displayed similar tolerance change when compared to the control groups (Figure 2G, 2I and S2E). Although pre-treatment with 0.1 mg/ml 2-AG for 2 days slightly decreased short-term ethanol tolerance, the difference was insignificant (Figure 2K). Likewise, the short-term ethanol tolerance change was not significantly different to the controls when the flies were pre-treated for 4 days (Figure S2G). Thus, these findings indicate that cannabinoids did not affect short-term tolerance of ethanol in flies.

**Figure 2.**
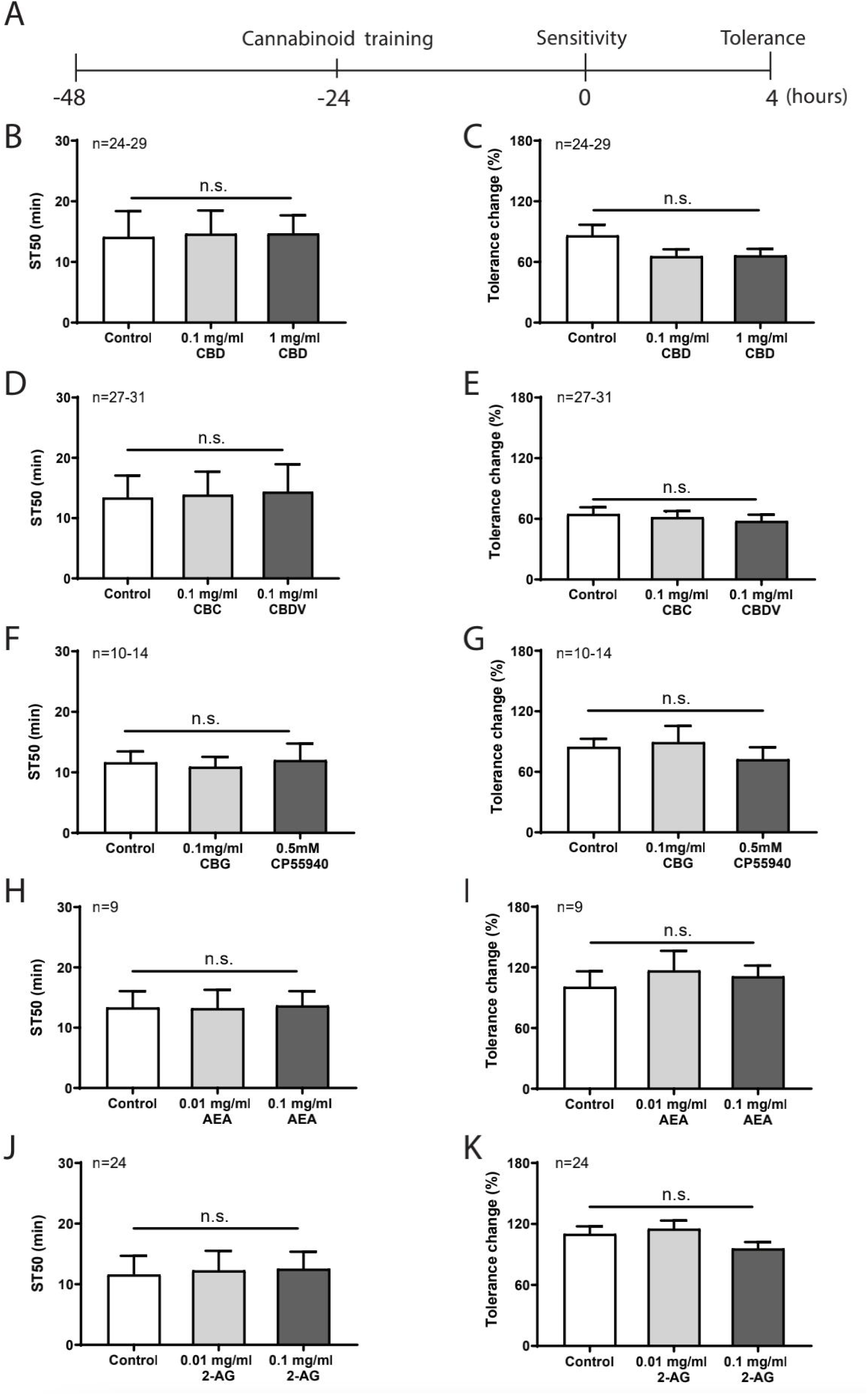
Cannabinoids did not alter short-term ethanol tolerance in flies. (A) A diagram of the experimental timeline for the ethanol sedation sensitivity and short-term ethanol tolerance assays. Flies were fed with various cannabinoids for 2 days before assessing the sedation sensitivity to ethanol at 0 hr and 4 hr. The change in ethanol sensitivity at both time points determines the short-term ethanol tolerance of the flies. Pre-treatment with phytocannabinoids including CBD at 0.1 mg/ml and 1 mg/ml (B, C), CBC, CBDV (D, E) and CBG (F, G) at 0.1 mg/ml, and synthetic cannabinoid CP55940 at 0.5 mM (F, G) did not alter both ethanol sedation sensitivity and short-term ethanol tolerance (n=10-29). (H, I) Flies pre-treated with AEA (H, I) and 2-AG (J, K) also did not affect both parameters (n=9-24). Statistical analysis was performed using one-way ANOVA. All data are represented as mean ± S.E.M.

### CBD and AEA reduced long-term tolerance to ethanol in flies

Following the assessment of short-term ethanol tolerance, we further characterize the effects of cannabinoids on long-term tolerance to ethanol. We focused on selective cannabinoids, namely CBD, CP55940, AEA and 2-AG, and investigated their potential effect on long-term ethanol tolerance. To this end, we first assessed innate ethanol sedation sensitivity on day 0 and repeated the same test once per day for 4 consecutive days (day 1 to 4) coupled with cannabinoid treatment (Figure 3A). Repeated exposure to 75% ethanol daily resulted in increased tolerance throughout the experiment from day 1 to day 4 (Figure 3B-E). Interestingly, flies treated with CBD at the low concentration of 0.1 mg/ml were less resistant to ethanol-induced sedation with the sedation time significantly lower from day 2 onwards when compared to the control-treated flies and this effect was sustained throughout the experiment (Figure 3B). However, CBD at the higher concentration of 1 mg/ml did not significantly suppress the ethanol tolerance although there is a trend towards decreased tolerance. There was a lack of CP55940 effect on long-term ethanol tolerance change (Figure 3C). Interestingly, treatment with AEA at both concentrations of 0.01 mg/ml and 0.1 mg/ml reduced sedation sensitivity to ethanol. However, the difference in the percentage of tolerance change was only significant at day 4 when flies were pre-treated with 0.1 mg/ml AEA (Figure 3D). However, treatment with 0.01 mg/ml and 0.1 mg/ml 2-AG did not affect long-term tolerance to ethanol (Figure 3E). These results indicate that CBD and AEA could mediate long-term tolerance to ethanol.

**Figure 3.**
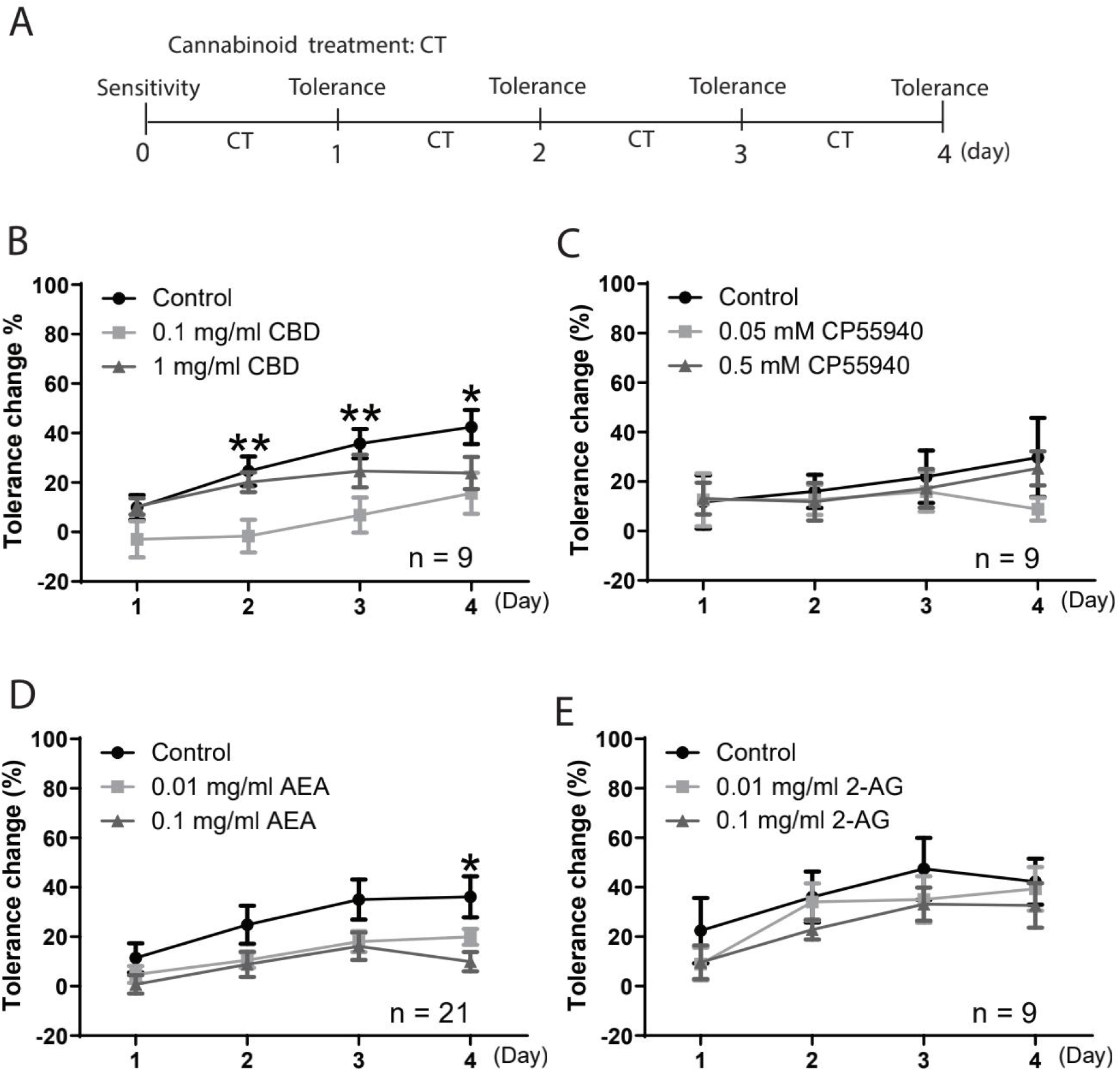
CBD and AEA reduced long-term tolerance to ethanol in flies. (A) A diagram of the experimental timeline of long-term ethanol tolerance assay in adult flies. Following the assessment of innate ethanol sedation sensitivity at day 0, the flies were treated with cannabinoids for 4 consecutive days and were subjected to ethanol sedation sensitivity assay once daily from day 1 to day 4. Treatment with 0.1 mg/ml CBD (B) and 0.1 mg/ml AEA (D) decreased the tolerance in ethanol sedation sensitivity from day 2 to day 4 (n=9-21). However, there was a lack of detectable effect on long-term ethanol tolerance with CP55940 (C) and 2-AG (E) pre-treatment (n=9). Statistical significance at each time point was determined using one-way ANOVA followed by Dunnett’s post-hoc test * p < 0.05 and ** p < 0.01. All data are represented as mean ± S.E.M.

### Cannabinoids did not alter ethanol-induced lethality in adult flies

Prolonged exposure to high concentrations of ethanol eventually resulted in sedation and subsequent death of the flies (Devineni and Heberlein, 2012). To explore whether cannabinoids could affect ethanol-induced lethality, the flies were pre-treated with cannabinoids for 2 or 4 consecutive days prior to exposure to high concentration of ethanol vapor at 100% (Figure 4A). Our initial findings in 3-day-old flies demonstrated that exposure to 100% ethanol for 35 minutes resulted in lethality rate of 12.73 ± 4.07 %. This was significantly increased to 42.50 ± 3.92% with 40 min ethanol exposure. With the extension of the duration of ethanol exposure to 43 and 45 minutes, the lethality rate of the flies was significantly higher at 59.44 ± 8.26 % and 66.36 ± 6.91%, respectively. These observations indicated a time-dependent effect on ethanol-induced lethality of the flies. Therefore, flies were subsequently exposed to 100% ethanol vapor for 45 minutes to understand the role of cannabinoids in ethanol-induced lethality. Pre-treatment of the flies with CBD and CP55940 for 2 days did not rescue or alleviate ethanol-induced lethality (Figure 4B). Likewise, flies fed with endocannabinoids of AEA (Figure 4D) and 2-AG (Figure 4F) for 2 days also resulted in high lethality which was similar to the flies fed with the respective controls. Prolonged 4-day pre-treatment with CBD, CP55940 (Figure 4C) and AEA (Figure 4E) also did not alter ethanol-induced lethality. Although the difference was insignificant, pre-treatment with 2-AG at the higher concentration of 0.1 mg/ml for 4 days displayed a slight reduction in the lethality of the flies (Figure 4G). Overall, our results suggest that cannabinoids play a negligible role in the modulation of ethanol-induced lethality in flies.

**Figure 4.**
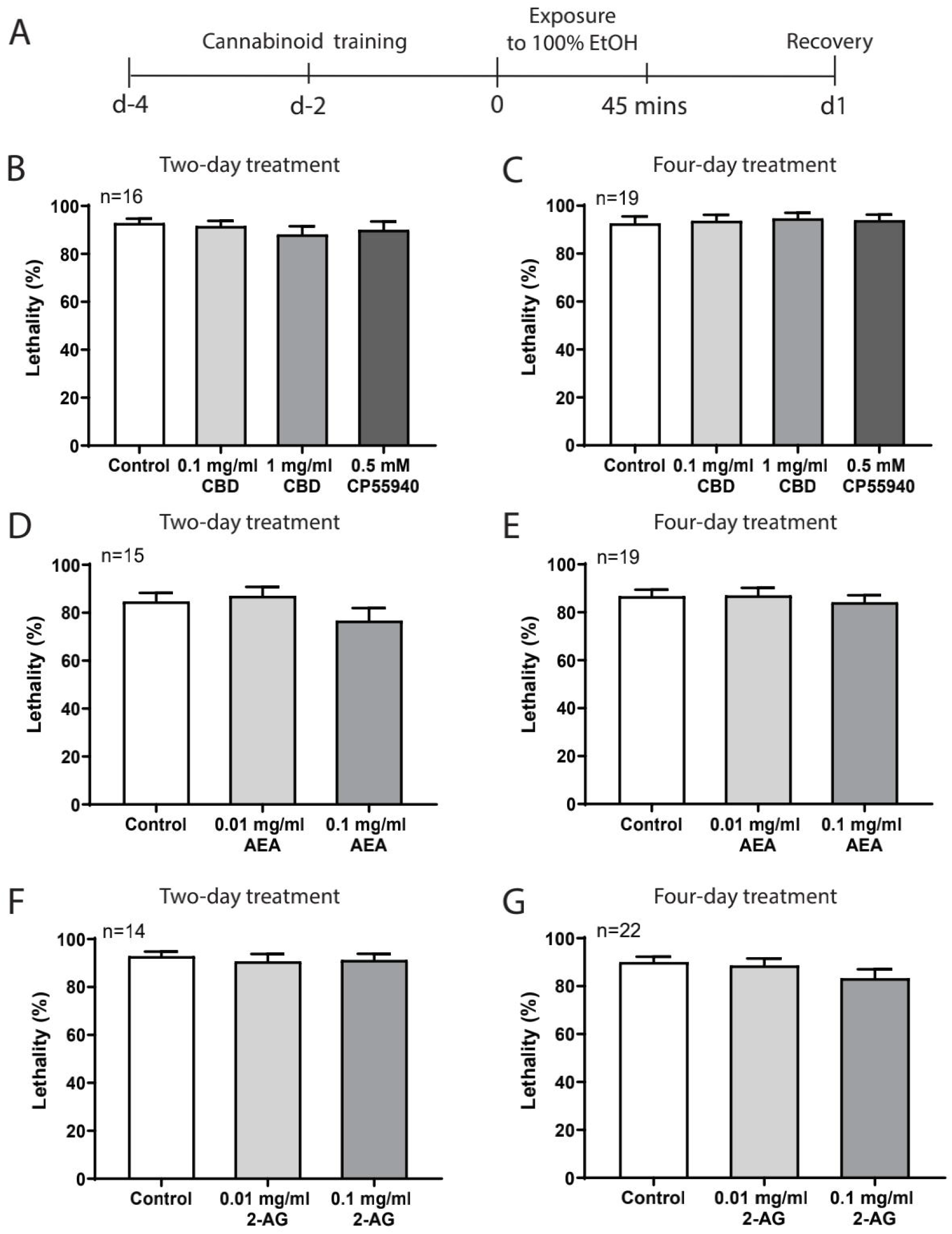
Cannabinoids did not alter ethanol-induced lethality in flies. (A) Flies were fed with cannabinoids for 2 (B, D, F) or 4 consecutive days (C, E, G) prior to exposure to 100% ethanol vapor for 45 minutes. The number of flies which died from ethanol exposure was examined 24 hours following recovery. Pre-treatment of CBD, CP55940 (B, C), AEA (D, E) and 2-AG (F, G) for 2 or 4 days displayed similar lethality percentage of flies after ethanol exposure (n=14-22). Statistical analysis was performed using one-way ANOVA. All data are represented as mean ± S.E.M.

### Cannabinoids did not rescue ethanol-induced developmental delay and lethality in *Drosophila*

Our observations of cannabinoid selectivity on various ethanol-induced behaviors prompted us to further study whether cannabinoids could affect larval/pupal development following ethanol exposure. It has been reported that exposure to ethanol during early developmental stages was detrimental to development and survival from embryogenesis to adult flies eclosion (McClure et al., 2011). This is consistent with our current findings that ethanol exposure to embryos at high concentrations of 7.5% and 10% significantly increased the number of days taken for pupal formation (Figure 5A). Eggs reared on non-ethanol food developed to pupae from 5.5 days after egg laying (AEL), with 50% of total pupae forming by 6.46 ± 0.13 days. The pupal formation was delayed with increasing ethanol concentrations from 5% to 10%. When exposed to 5% and 7.5% ethanol, the respective average time taken for formation of 50% pupae (DT50) was 7.29 ± 0.24 days and 8.19 ± 0.33 days. As compared to the non-ethanol conditions, there was an approximate delay of 4 days for pupal formation at 10.46 ± 0.22 days with 10% ethanol exposure. While increasing ethanol concentrations delayed pupal development, there was a significant ∼50% decrease in the relative number of pupae forming with 10% ethanol exposure as compared to the control group (Figure 5B). This indicates that ethanol-reared embryos resulted in larval developmental delay, retarded pupal formation and even lethality. Moreover, ethanol exposure also exhibited a dose-dependent effect on the size of larvae and pupae. The average size of the control 3^rd^ instar wandering larvae (wL3) and pupae was 18.03 ± 0.37 mm^2^ and 14.68 ± 0.36 mm^2^, respectively. When the eggs were reared on 10% ethanol containing food medium, there was a ∼30% increase in the size of larvae to 23.59 ± 0.43 mm^2^ and pupae to 19.03 ± 0.28 mm^2^. There is a negative correlation of increasing ethanol levels with larval development/survival, and a positive relationship with larval/pupal body size.

**Figure 5.**
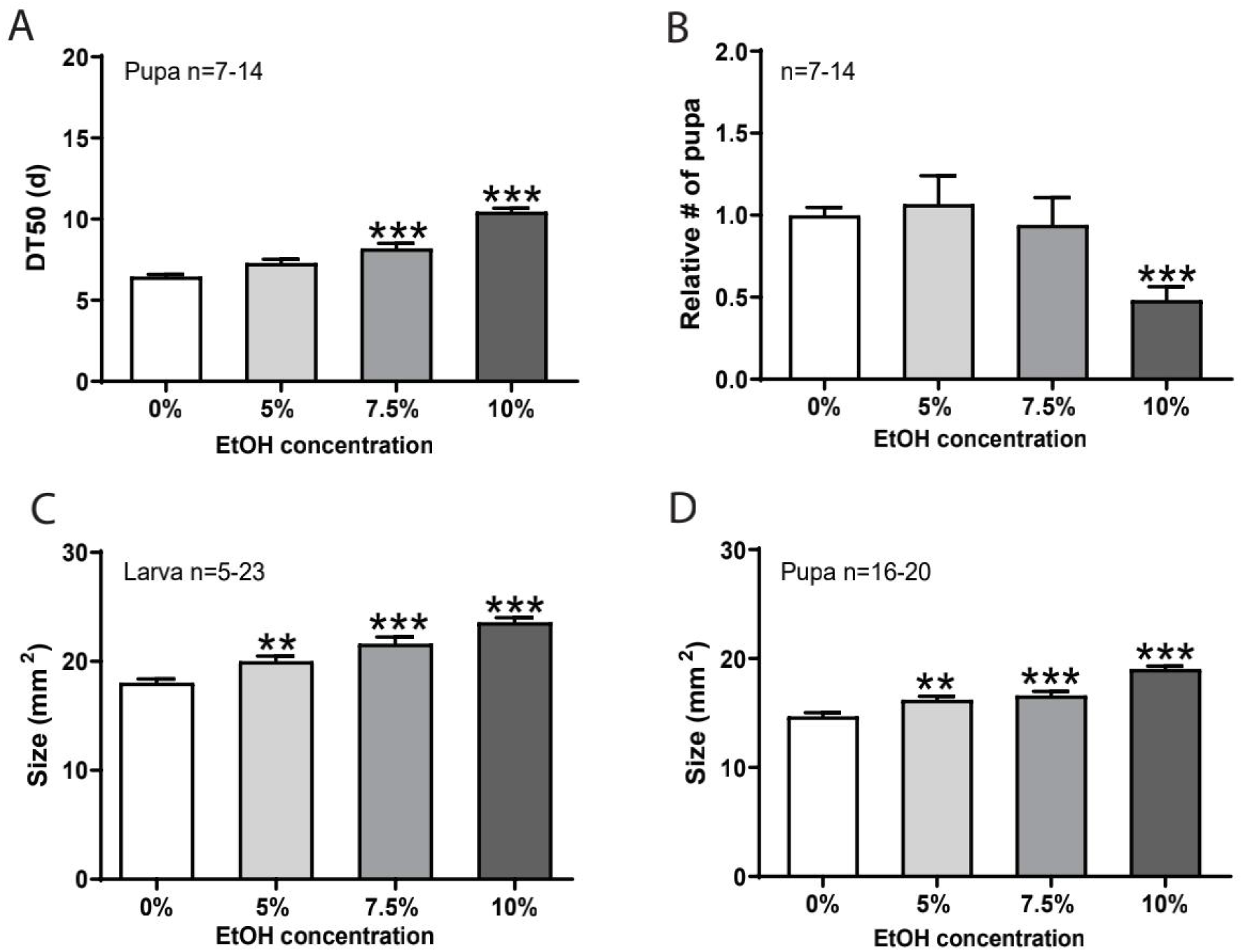
Exposure to ethanol delayed larval development and increased larval/pupal size. Eggs were collected and reared on normal food medium containing ethanol with increasing concentrations from 0% to 10%. (A) Exposure to high concentrations of ethanol at 7.5% and 10% significantly increased the time taken for the development of 50% eggs to pupae (DT50) (n=7-14). (B) The relative number of pupae formed was significantly lower as compared to the control group when exposed to 10% ethanol only (n=7-14). Ethanol exposure increased the size of larvae (C) and pupae (D) in a dose-dependent manner (n=5-23). Statistical significance was performed using one-way ANOVA followed by Dunnett’s post-hoc test, ** p < 0.01 and *** p < 0.001. Data are represented as mean ± S.E.M.

A recent study has shown that treatment of CBD significantly improved the survival rate of ethanol-intoxicated *M. sexta* larvae (Park et al., 2019). We next determined whether cannabinoids could rescue or alleviate ethanol-induced developmental delay and/or lethality. Eggs raised on food medium containing 10% ethanol and CBD at 0.02 mg/ml or 0.1 mg/ml displayed slightly longer period of larval development and lower survival rate when compared to the control group raised on food medium containing 10% ethanol only (Figure 6A and 6B). Despite that, exposure of the eggs to CBD alone appeared to display adverse effects on the development of eggs to pupae in a dose-dependent manner (Figure S3A). Furthermore, treatment with CP55940 only (Figure S3B) or CP55940 with 10% ethanol (Figure 6C-D) did not affect larval development and viability. Unexpectedly, ethanol-induced developmental delay was significantly deteriorated upon 0.01 mg/ml AEA (Figure 6E) or 2-AG co-treatment (Figure 6G). Despite a trend towards decreasing viability with endocannabinoid treatments, the relative number of pupae was not significantly different from the control groups (Figure 6F and 6H). When the eggs were reared on food medium containing AEA or 2-AG alone, the time taken for pupal formation was unaltered (Figure S3C-D). In contrast to the previous study that CBD treatment alleviates ethanol-induced death in the tobacco hornworm (Park et al., 2019), our findings unexpectedly suggest that the detrimental effects of ethanol-induced developmental delay and toxicity are not attenuated by CBD or CP55940 treatment, instead, are further exacerbated by endocannabinoids.

**Figure 6.**
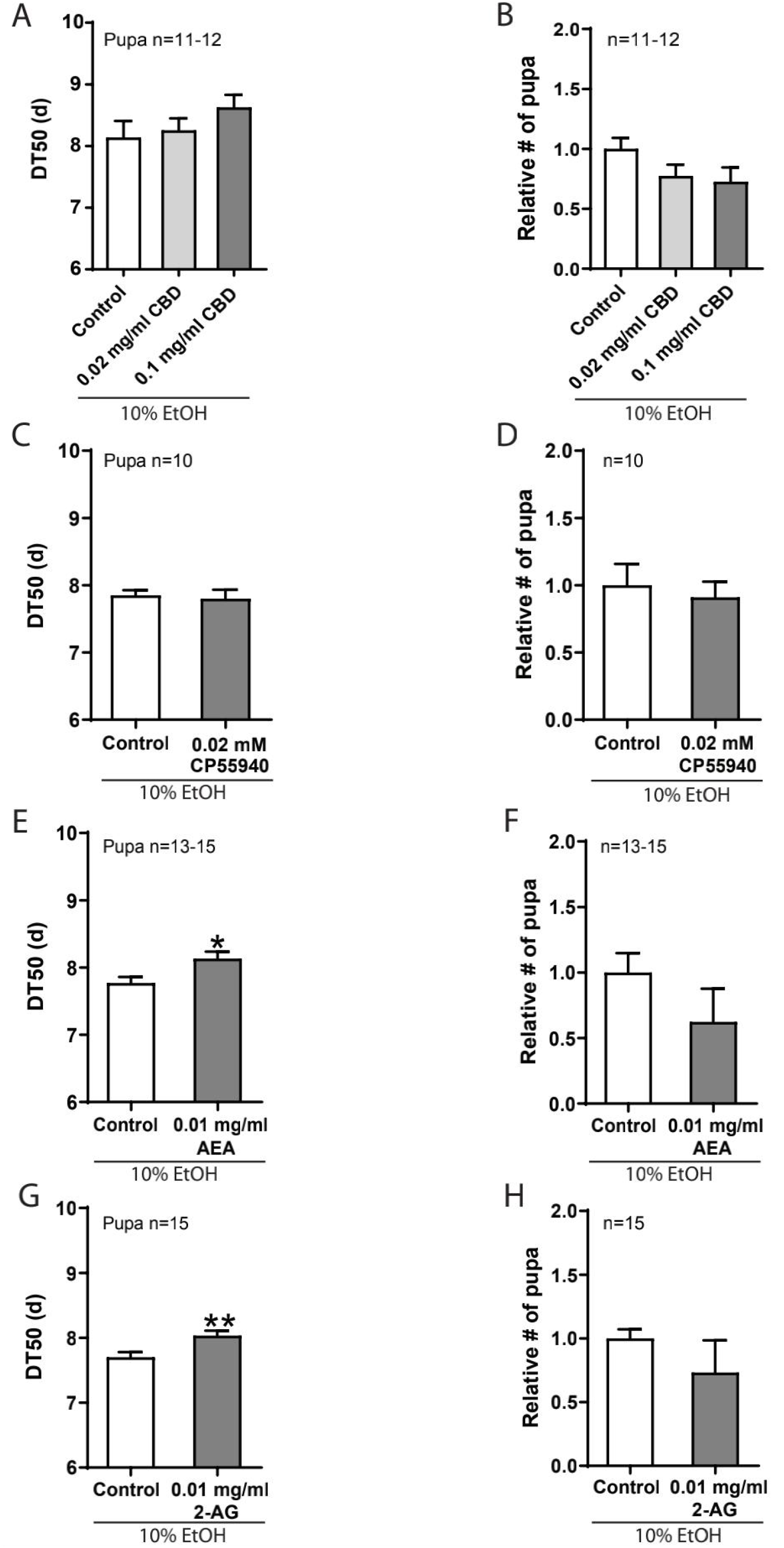
Cannabinoids did not rescue ethanol-induced developmental delay and lethality in *Drosophila*. Eggs were raised on food medium with 10% ethanol and cannabinoids or the respective control solutions. The time taken for the development from eggs to pupae (left panel) and survival of pupae (right panel) were assessed in this assay. There was no effect of CBD (A, B) and CP55940 (C, D) on the development and survival of egg-to-pupa (n=10-12). However, treatment with AEA (E) or 2-AG (G) significantly increased time taken for pupal formation (n=13-15). Although AEA (F) and 2-AG (H) decreased ethanol-induced lethality, the difference was not significant. Statistical analysis was performed using two-tailed unpaired t-test or one-way ANOVA followed by Dunnett’s post-hoc test (n=10-15 vials/group). All data are represented as mean ± S.E.M.

## Discussion

Previous studies using the *Drosophila* model have highlighted the protective role of cannabinoids in food intake (He et al., 2021), seizure (Jacobs and Sehgal, 2020), paraquat-evoked toxicity (Jimenez-Del-Rio et al., 2008) and cardiac function (Gómez et al., 2019). However, it remains unexplored whether cannabinoids affect ethanol responses in *Drosophila* which has been shown to be effective in modelling complex ethanol behaviors in mammals. It is well established that adult flies displayed preference to ethanol, resistance to ethanol sedation and lethality following prolonged ethanol exposure (Devineni and Heberlein, 2009; Devineni and Heberlein, 2012; Scholz et al., 2000). Furthermore, early exposure to ethanol induced developmental delay (McClure et al., 2011). In this current study, we provide new insights into the role of cannabinoids in ethanol-related behaviors. We systematically investigated the effects of cannabinoids on a spectrum of behaviors including ethanol preference, ethanol sedation sensitivity and ethanol lethality, and ethanol-induced developmental delay in *Drosophila*.

Amongst the various cannabinoids, the non-psychoactive phytocannabinoid CBD and endocannabinoids exhibited selectivity on ethanol behaviors with the observations of decrease in preferential ethanol consumption or long-term ethanol tolerance. These results appear to be promising as reduction in ethanol intake and tolerance could possibly decrease the likelihood of progression into chronic ethanol-related health issues. Similar to our findings, a recent study has reported that treatment with CBD in mice reduced ethanol preference and consumption in a two-bottle choice paradigm in a dose-dependent manner (Viudez-Martínez et al., 2018). It is evident that ethanol preference and ethanol tolerance are modulated by the canonical endocannabinoid system whereby the biological effects of the two main endocannabinoids, AEA and 2-AG, are mediated mainly through the CB_1/2_ receptors (Cristino et al., 2020). In contrast to our findings, many studies in the rodent models have shown opposing effects of endocannabinoids on ethanol preference. High ethanol preference in mice was significantly decreased with blockade of the canonical CB_1/2_ receptors or in CB_1_ receptor knock out mice (Gallate and McGregor, 1999; Hungund et al., 2003; Wang et al., 2003). Furthermore, *in vitro* studies demonstrated that chronic exposure to ethanol led to accumulation of AEA (Basavarajappa and Hungund, 1999a; Basavarajappa et al., 2003) and 2-AG (Basavarajappa et al., 2000). The upregulation of endocannabinoid levels led to persistent activation of CB_1/2_ receptors, thus resulting in the downregulation of the availability of these receptors (Basavarajappa and Hungund, 1999b). Both endocannabinoids, AEA and 2-AG, are inactivated through intracellular enzymatic degradation through fatty acid amidohydrolase (FAAH)-mediated hydrolysis to arachidonic acid (AA) and ethanolamine or glycerol, respectively (Basavarajappa, 2007). With chronic exposure to ethanol, the impairment of degradation machinery could account for the accumulation of endocannabinoids. Previously, the findings of *in vivo* studies reported increase in ethanol preference and decrease in ethanol sensitivity in mice administered with FAAH inhibitors and FAAH knock out mice (Blednov et al., 2007; Falenski et al., 2010), indicative of the importance of endocannabinoids in the regulation of ethanol preference and sedation. The contrasting effects of cannabinoids in different animal models could be due to the diversity of the cannabinoid system. It is evident that the canonical CB_1/2_ receptor-mediated signaling pathway is non-existent in *Drosophila* due to the absence of these receptors (McPartland et al., 2001). We thereby postulate that endocannabinoids could exert their effects via identified non-canonical receptors or unknown CB_1/2_ like receptors. Notably, CBD exhibits low binding affinity to the canonical receptors (Cristino et al., 2020) and is known to have diverse interaction with other systems through various GPCRs including GPR55, 5HT_1A_ receptors, TRPV1 channels and opioid receptors (De Petrocellis et al., 2011; Kathmann et al., 2006; Russo et al., 2005; Ryberg et al., 2007). The selectivity of CBD in the ethanol behaviors could be modulated through different GPCR-mediated pathways. However, more work is required to unravel the underlying mechanisms of these cannabinoids in ethanol behaviors. Apart from the possibility of differential modulation of the cannabinoids on these ethanol behaviors via different pathways, the processes mediating these ethanol behaviors are dissociable in *Drosophila*. For instance, the male flies were found to be more susceptible to sedation and less resistant to ethanol lethality (Devineni and Heberlein, 2012). Our findings demonstrated that treatment with CBD and endocannabinoid AEA suppressed long-term ethanol tolerance without affecting ethanol-induced lethality, suggesting that these cannabinoids have specific functional implications in chronic ethanol responses.

As selective cannabinoids displayed protective effects on ethanol preference and long-term ethanol tolerance in the adult flies, it will be interesting to determine whether cannabinoids affect development in *Drosophila*. Exposure to CBD alone, but not CP55940 and endocannabinoids, led to a significant delay in normal larval development in *Drosophila*. This finding is consistent with a study in tobacco hornworm *Manduca sexta* whereby the larvae reared on food containing high concentration of CBD displayed significant reduction in size and increased lethality (Park et al., 2019). Exposure to CP55940 throughout the neurulation stage, a critical morphogenetic event during human gestation, induced craniofacial and brain abnormalities in mice (Fish et al., 2016; Gilbert et al., 2016) and zebrafish (Fish et al., 2019). The naturally-occurring CBD and synthetic cannabinoid CP55940 have adverse effects on the processes governing normal development possibly due to disruptions to cannabinoid-signaling pathway during growth. The innate endocannabinoid system plays a critical role in the regulation of fundamental developmental processes, especially development of the central nervous system and synaptic communication (Basavarajappa et al., 2009; Clarke et al., 2021; Harkany et al., 2007).

As the combinatorial intake of marijuana/*Cannabis* and alcohol is prominently increasing in pregnant women (Young-Wolff et al., 2017), we aim to understand the relationship of ethanol and cannabinoids on development in *Drosophila*. The interesting and unexpected finding in our study is that endocannabinoids exacerbated ethanol-induced embryo-to-pupal developmental delay. Although exogenous administration of endocannabinoids alone did not affect normal development, combinatorial exposure to these cannabinoids and ethanol negatively impacted development and viability in *Drosophila*. Based on these observations, endocannabinoids function differently in the absence or presence of ethanol, suggesting that a potential interaction between endocannabinoids and ethanol is detrimental to development. Endocannabinoids have been shown to be conserved inhibitors of Hedgehog signaling (Khaliullina et al., 2015). A recent study has demonstrated that cannabinoids exacerbated alcohol teratogenesis through CB_1_ receptor-Hedgehog pathway in zebrafish (Fish et al., 2019). *In vivo* studies using the rodent models have revealed that postnatal exposure to ethanol increased the levels of AEA (Vinod et al., 2006) but not 2-AG and CB_1_ receptors, thus leading to neurodegeneration through the AEA/CB_1_ receptor signaling pathway (Subbanna et al., 2013). It was recently shown that AEA analog, Arachidonoyl-20-chloroethylamide, with low amount of ethanol induced dysmorphogenesis in zebrafish (Boa-Amponsem et al., 2019). In addition, we also observe a slight delay in ethanol-induced pupal development and decrease in viability with CBD treatment although the difference was insignificant. In line with our finding, prenatal exposure to CBD enhanced the eye and face malformation induced by ethanol (Fish et al., 2019). However, CBD was found to be a rescuing agent following exposure to ethanol in the tobacco hornworm (Park et al., 2019). Notably, other cannabinoids including the psychoactive Δ^9^-THC, synthetic cannabinoids HU-210 and CP 55,940 also worsened ethanol-induced birth defects (Fish et al., 2019) and neurodegeneration (Hansen et al., 2008).

The differential effects of cannabinoids in different ethanol responses could be due to several reasons. Firstly, the endocannabinoid system differs between the model organisms. It is notable that the canonical CB_1/2_ receptors are absent in *Drosophila* while being abundant in mammals (McPartland et al., 2001). The mechanistic pathway of the cannabinoids in the modulation of ethanol behaviors in *Drosophila* is presumably mediated via non-CB_1/2_ receptors. The identified non-canonical receptors of CBD and endocannabinoids are 5-HT_1A_ receptors, GPR55, TRPV1 channels, opioid receptors and peroxisome proliferator-activated receptor (PPAR) (Cristino et al., 2020). Secondly, the selectivity of cannabinoids could possibly to due to the exposure of the cannabinoids at different developmental stages. The cannabinoids could function differently on the developing and mature systems during development and adulthood.

This present study has characterized the effects of cannabinoids on various ethanol-related responses in *Drosophila*. Our results demonstrate that selective phytocannabinoid and endocannabinoids reduced ethanol preference and long-term ethanol tolerance in adult flies. Despite being protective towards selective ethanol-related behaviors, these cannabinoids function differently in the presence of ethanol during the developmental stages. Combinatorial exposure to ethanol and endocannabinoids is detrimental to larval development. In conclusion, this study has shown evidence of differential modulation of cannabinoids on ethanol behaviors.

## Figure legends

**Figure S1.**
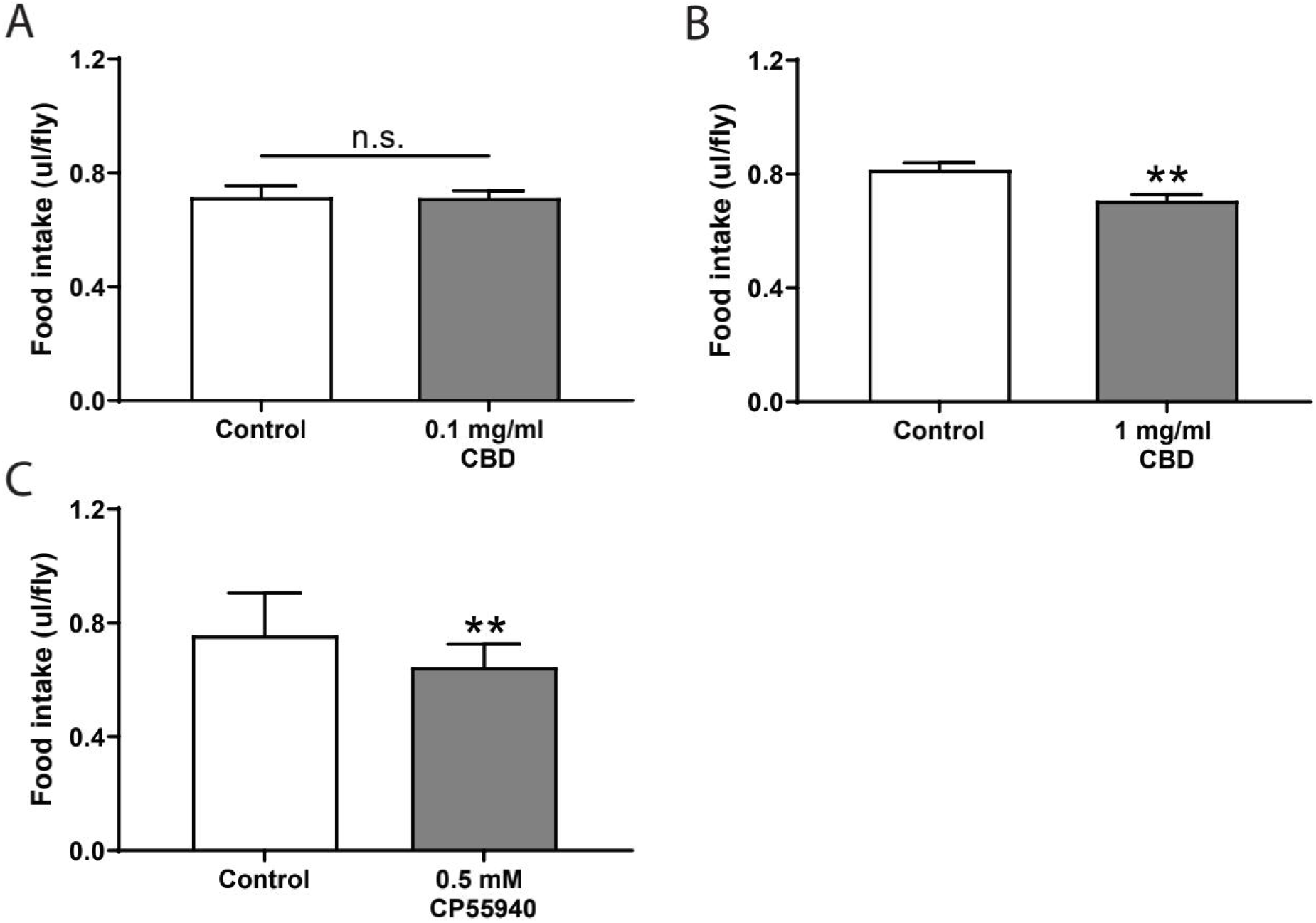
CBD and CP55940 treatments reduced food intake in flies. Following 2-day pre-treatment of cannabinoids, the flies were presented with non-ethanol or ethanol containing liquid food for 2 consecutive days. The average daily food consumption following cannabinoid treatment was recorded. Pre-treatment with 1 mg/ml CBD (B), but not 0.1 mg/ml CBD (A) significantly inhibited food intake. Flies pre-treated with 0.5 mM CP55940 (C) consumed significantly less food as compared to the control groups. Statistical significance was determined using one-way ANOVA followed by Dunnett’s post-hoc test, * p < 0.05, ** p < 0.01 and *** p < 0.001 (n=18-22 vials/group). The data are represented as mean ± S.E.M.

**Figure S2.**
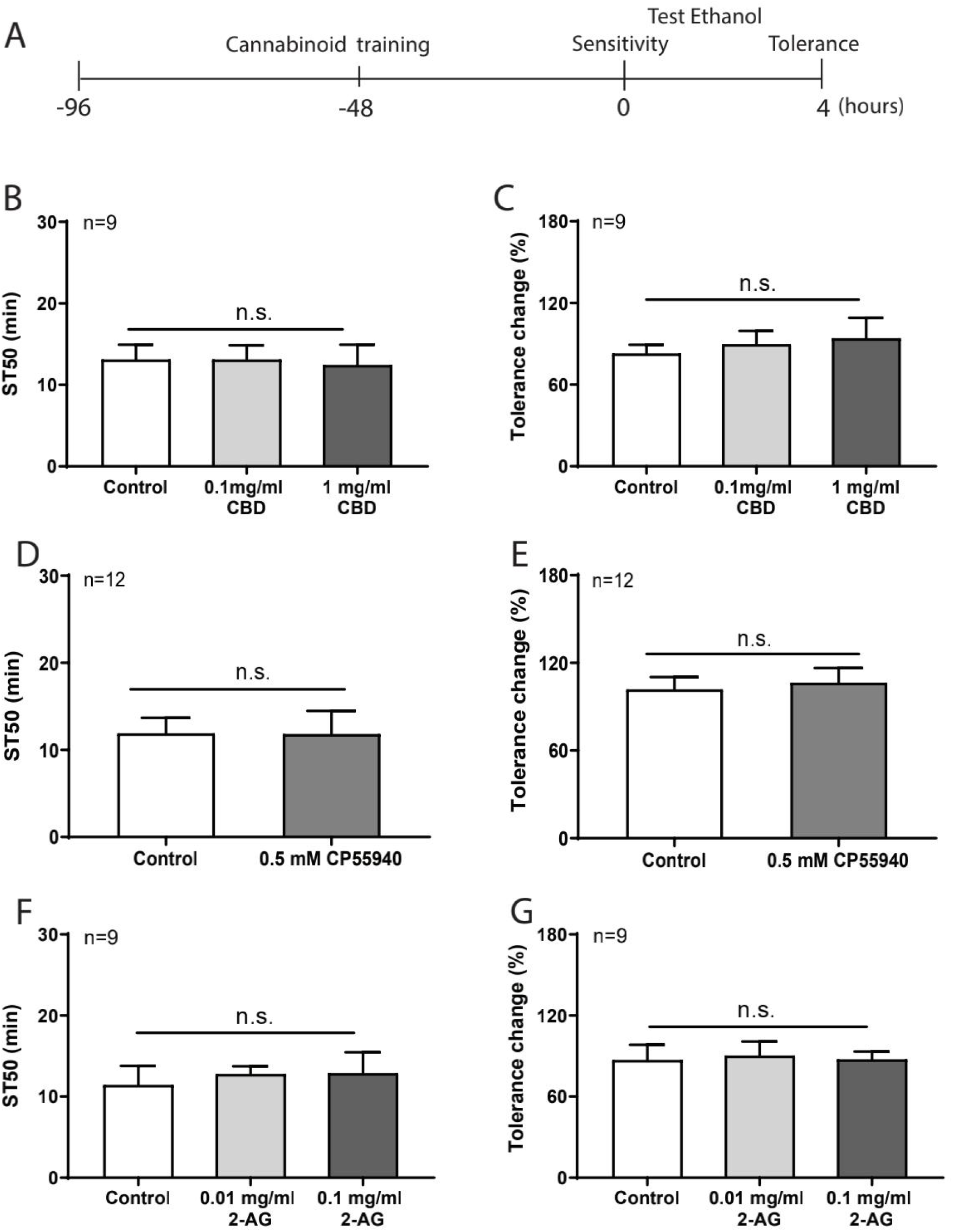
4-day pre-treatment of cannabinoids did not affect ethanol sedation sensitivity and short-term ethanol tolerance. (A) The experimental timeline for the ethanol sedation sensitivity and short-term ethanol tolerance assay where the flies were pre-treated with cannabinoids for 4 consecutive days before the behavioral test. Pre-treatment with CBD at 0.1 mg/ml and 1 mg/ml (B, C), CP55940 at 0.5 mM (D, E) and 2-AG at 0.01 mg/ml and 0.1 mg/ml (F, G) did not affect sedation sensitivity and short-term tolerance to ethanol (n=9-12). Statistical analysis was performed using one-way ANOVA. All data are represented as mean ± S.E.M.

**Figure S3.**
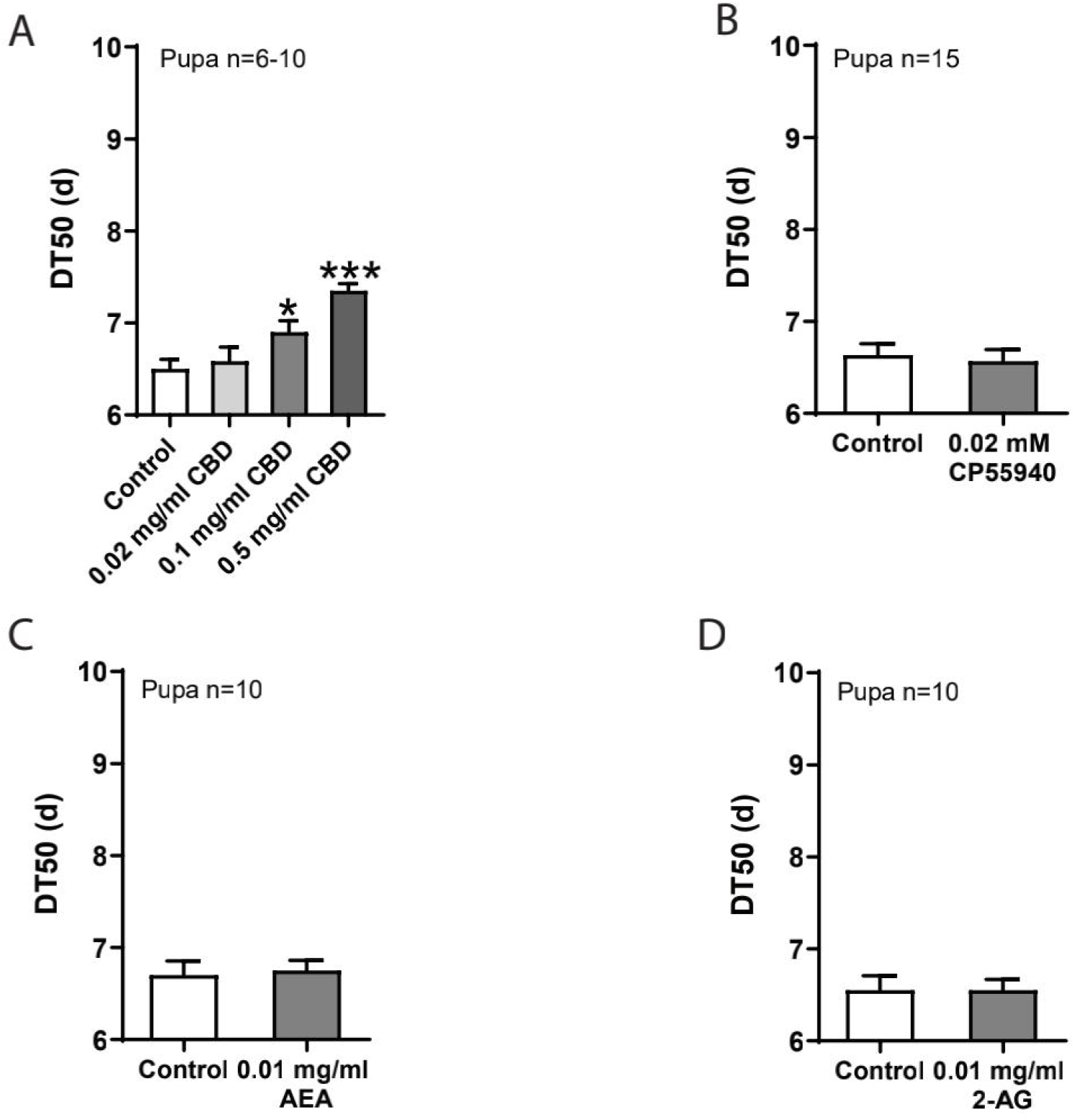
CBD delayed larval development. Eggs were raised on cannabinoid-containing food medium. (A, B) CBD at 0.1 mg/ml and 0.5 mg/ml, but not at 0.02 mg/ml, significantly delayed the larval development (A) (n=6-10). Treatment with CP55940 (B), AEA (C) and 2-AG (D) did not affect normal pupal formation (n=10-15). One-way ANOVA followed by Dunnett’s post-hoc test was applied to determine statistical significance, *** p < 0.001.

## References

Bainton, R.J., Tsai, L.T., Singh, C.M., Moore, M.S., Neckameyer, W.S., and Heberlein, U. (2000). Dopamine modulates acute responses to cocaine, nicotine and ethanol in Drosophila. Current biology : CB 10, 187–194.

Basavarajappa, B.S. (2007). Critical enzymes involved in endocannabinoid metabolism. Protein and peptide letters 14, 237–246.

Basavarajappa, B.S., and Hungund, B.L. (1999a). Chronic ethanol increases the cannabinoid receptor agonist anandamide and its precursor N-arachidonoylphosphatidylethanolamine in SK-N-SH cells. Journal of neurochemistry 72, 522–528.

Basavarajappa, B.S., and Hungund, B.L. (1999b). Down-regulation of cannabinoid receptor agonist-stimulated [35S]GTP gamma S binding in synaptic plasma membrane from chronic ethanol exposed mouse. Brain research 815, 89–97.

Basavarajappa, B.S., Nixon, R.A., and Arancio, O. (2009). Endocannabinoid system: emerging role from neurodevelopment to neurodegeneration. Mini Rev Med Chem 9, 448–462.

Basavarajappa, B.S., Saito, M., Cooper, T.B., and Hungund, B.L. (2000). Stimulation of cannabinoid receptor agonist 2-arachidonylglycerol by chronic ethanol and its modulation by specific neuromodulators in cerebellar granule neurons. Biochimica et Biophysica Acta (BBA) - Molecular Basis of Disease 1535, 78–86.

Basavarajappa, B.S., Saito, M., Cooper, T.B., and Hungund, B.L. (2003). Chronic ethanol inhibits the anandamide transport and increases extracellular anandamide levels in cerebellar granule neurons. European journal of pharmacology 466, 73–83.

Berger, K.H., Kong, E.C., Dubnau, J., Tully, T., Moore, M.S., and Heberlein, U. (2008). Ethanol sensitivity and tolerance in long-term memory mutants of Drosophila melanogaster. Alcoholism, clinical and experimental research 32, 895–908.

Blednov, Y.A., Cravatt, B.F., Boehm, S.L., Walker, D., and Harris, R.A. (2007). Role of Endocannabinoids in Alcohol Consumption and Intoxication: Studies of Mice Lacking Fatty Acid Amide Hydrolase. Neuropsychopharmacology 32, 1570–1582.

Boa-Amponsem, O., Zhang, C., Mukhopadhyay, S., Ardrey, I., and Cole, G.J. (2019). Ethanol and cannabinoids interact to alter behavior in a zebrafish fetal alcohol spectrum disorder model. Birth defects research 111, 775–788.

Carvalho, A.F., Heilig, M., Perez, A., Probst, C., and Rehm, J. (2019). Alcohol use disorders. The Lancet 394, 781–792.

Clarke, T.L., Johnson, R.L., Simone, J.J., and Carlone, R.L. (2021). The Endocannabinoid System and Invertebrate Neurodevelopment and Regeneration. International Journal of Molecular Sciences 22.

Cristino, L., Bisogno, T., and Di Marzo, V. (2020). Cannabinoids and the expanded endocannabinoid system in neurological disorders. Nature reviews Neurology 16, 9–29.

De Petrocellis, L., Ligresti, A., Moriello, A.S., Allarà, M., Bisogno, T., Petrosino, S., Stott, C.G., and Di Marzo, V. (2011). Effects of cannabinoids and cannabinoid-enriched Cannabis extracts on TRP channels and endocannabinoid metabolic enzymes. British journal of pharmacology 163, 1479–1494.

De Ternay, J., Naassila, M., Nourredine, M., Louvet, A., Bailly, F., Sescousse, G., Maurage, P., Cottencin, O., Carrieri, P.M., and Rolland, B. (2019). Therapeutic Prospects of Cannabidiol for Alcohol Use Disorder and Alcohol-Related Damages on the Liver and the Brain. Front Pharmacol 10, 627–627.

Devineni, A.V., and Heberlein, U. (2009). Preferential ethanol consumption in Drosophila models features of addiction. Current biology : CB 19, 2126–2132.

Devineni, A.V., and Heberlein, U. (2012). Acute ethanol responses in Drosophila are sexually dimorphic. Proceedings of the National Academy of Sciences 109, 21087.

Ehrhart, F., Roozen, S., Verbeek, J., Koek, G., Kok, G., van Kranen, H., Evelo, C.T., and Curfs, L.M.G. (2019). Review and gap analysis: molecular pathways leading to fetal alcohol spectrum disorders. Molecular Psychiatry 24, 10–17.

ElSohly, M., and Gul, W. (2015). Constituents of Cannabit Sativa. Handbook of Cannabis.

Falenski, K.W., Thorpe, A.J., Schlosburg, J.E., Cravatt, B.F., Abdullah, R.A., Smith, T.H., Selley, D.E., Lichtman, A.H., and Sim-Selley, L.J. (2010). FAAH−/− Mice Display Differential Tolerance, Dependence, and Cannabinoid Receptor Adaptation After Δ9-Tetrahydrocannabinol and Anandamide Administration. Neuropsychopharmacology 35, 1775–1787.

Fish, E.W., Holloway, H.T., Rumple, A., Baker, L.K., Wieczorek, L.A., Moy, S.S., Paniagua, B., and Parnell, S.E. (2016). Acute alcohol exposure during neurulation: Behavioral and brain structural consequences in adolescent C57BL/6J mice. Behav Brain Res 311, 70–80.

Fish, E.W., Murdaugh, L.B., Zhang, C., Boschen, K.E., Boa-Amponsem, O., Mendoza-Romero, H.N., Tarpley, M., Chdid, L., Mukhopadhyay, S., Cole, G.J., et al. (2019). Cannabinoids Exacerbate Alcohol Teratogenesis by a CB1-Hedgehog Interaction. Scientific reports 9, 16057–16057.

Gallate, J.E., and McGregor, I.S. (1999). The motivation for beer in rats: effects of ritanserin, naloxone and SR 141716. Psychopharmacology 142, 302–308.

Gilbert, M.T., Sulik, K.K., Fish, E.W., Baker, L.K., Dehart, D.B., and Parnell, S.E. (2016). Dose-dependent teratogenicity of the synthetic cannabinoid CP-55,940 in mice. Neurotoxicology and Teratology 58, 15–22.

Gómez, I.M., Rodríguez, M.A., Santalla, M., Kassis, G., Colman Lerner, J.E., Aranda, J.O., Sedán, D., Andrinolo, D., Valverde, C.A., and Ferrero, P. (2019). Inhalation of marijuana affects Drosophila heart function. Biology Open 8, bio044081.

Hansen, H.H., Krutz, B., Sifringer, M., Stefovska, V., Bittigau, P., Pragst, F., Marsicano, G., Lutz, B., and Ikonomidou, C. (2008). Cannabinoids enhance susceptibility of immature brain to ethanol neurotoxicity. Annals of neurology 64, 42–52.

Hanuš, L.O., Meyer, S.M., Muñoz, E., Taglialatela-Scafati, O., and Appendino, G. (2016). Phytocannabinoids: a unified critical inventory. Natural product reports 33, 1357–1392.

Harkany, T., Guzmán, M., Galve-Roperh, I., Berghuis, P., Devi, L.A., and Mackie, K. (2007). The emerging functions of endocannabinoid signaling during CNS development. Trends in Pharmacological Sciences 28, 83–92.

He, J., Tan, A.M.X., Ng, S.Y., Rui, M., and Yu, F. (2021). Cannabinoids modulate food preference and consumption in Drosophila melanogaster. Scientific Reports 11, 4709.

Hungund, B.L., Szakall, I., Adam, A., Basavarajappa, B.S., and Vadasz, C. (2003). Cannabinoid CB1 receptor knockout mice exhibit markedly reduced voluntary alcohol consumption and lack alcohol-induced dopamine release in the nucleus accumbens. Journal of neurochemistry 84, 698–704.

Ja, W.W., Carvalho, G.B., Mak, E.M., de la Rosa, N.N., Fang, A.Y., Liong, J.C., Brummel, T., and Benzer, S. (2007). Prandiology of Drosophila and the CAFE assay. Proc Natl Acad Sci U S A 104, 8253–8256.

Jacobs, J.A., and Sehgal, A. (2020). Anandamide Metabolites Protect against Seizures through the TRP Channel Water Witch in Drosophila melanogaster. Cell reports 31, 107710.

Jimenez-Del-Rio, M., Daza-Restrepo, A., and Velez-Pardo, C. (2008). The cannabinoid CP55,940 prolongs survival and improves locomotor activity in Drosophila melanogaster against paraquat: implications in Parkinson’s disease. Neuroscience research 61, 404–411.

Kathmann, M., Flau, K., Redmer, A., Tränkle, C., and Schlicker, E. (2006). Cannabidiol is an allosteric modulator at mu-and delta-opioid receptors. Naunyn-Schmiedeberg’s archives of pharmacology 372, 354–361.

Khaliullina, H., Bilgin, M., Sampaio, J.L., Shevchenko, A., and Eaton, S. (2015). Endocannabinoids are conserved inhibitors of the Hedgehog pathway. Proc Natl Acad Sci U S A 112, 3415–3420.

Koren, G., Cohen, R., and Sachs, O. (2021). Use of Cannabis in Fetal Alcohol Spectrum Disorder. Cannabis and cannabinoid research 6, 74–76.

Lange, S., Probst, C., Gmel, G., Rehm, J., Burd, L., and Popova, S. (2017). Global Prevalence of Fetal Alcohol Spectrum Disorder Among Children and Youth: A Systematic Review and Meta-analysis. JAMA Pediatrics 171, 948–956.

Maples, T., and Rothenfluh, A. (2011). A simple way to measure ethanol sensitivity in flies. J Vis Exp, 2541.

Marquardt, K., and Brigman, J.L. (2016). The impact of prenatal alcohol exposure on social, cognitive and affective behavioral domains: Insights from rodent models. Alcohol 51, 1–15.

McClure, K.D., French, R.L., and Heberlein, U. (2011). A Drosophila model for fetal alcohol syndrome disorders: role for the insulin pathway. Disease Models & Mechanisms 4, 335.

McKenzie, J.A., and McKechnie, S.W. (1979). A comparative study of resource utilization in natural populations of Drosophila melanogaster and D. simulans. Oecologia 40, 299–309.

McPartland, J., Di Marzo, V., De Petrocellis, L., Mercer, A., and Glass, M. (2001). Cannabinoid receptors are absent in insects. The Journal of comparative neurology 436, 423–429.

Mechoulam, R., Hanuš, L.O., Pertwee, R., and Howlett, A.C. (2014). Early phytocannabinoid chemistry to endocannabinoids and beyond. Nature Reviews Neuroscience 15, 757–764.

Moore, M.S., DeZazzo, J., Luk, A.Y., Tully, T., Singh, C.M., and Heberlein, U. (1998). Ethanol Intoxication in Drosophila: Genetic and Pharmacological Evidence for Regulation by the cAMP Signaling Pathway. Cell 93, 997–1007.

Park, S.-H., Staples, S.K., Gostin, E.L., Smith, J.P., Vigil, J.J., Seifried, D., Kinney, C., Pauli, C.S., and Heuvel, B.D.V. (2019). Contrasting Roles of Cannabidiol as an Insecticide and Rescuing Agent for Ethanol–induced Death in the Tobacco Hornworm Manduca sexta. Scientific Reports 9, 10481.

Park, S.J., and Ja, W.W. (2020). Absolute ethanol intake predicts ethanol preference in Drosophila melanogaster. The Journal of Experimental Biology 223, jeb224121.

Parkhurst, S.J., Adhikari, P., Navarrete, J.S., Legendre, A., Manansala, M., and Wolf, F.W. (2018). Perineurial Barrier Glia Physically Respond to Alcohol in an Akap200-Dependent Manner to Promote Tolerance. Cell reports 22, 1647–1656.

Petruccelli, E., Li, Q., Rao, Y., and Kitamoto, T. (2016). The Unique Dopamine/Ecdysteroid Receptor Modulates Ethanol-Induced Sedation in Drosophila. The Journal of neuroscience : the official journal of the Society for Neuroscience 36, 4647–4657.

Russo, E.B., Burnett, A., Hall, B., and Parker, K.K. (2005). Agonistic properties of cannabidiol at 5-HT1a receptors. Neurochemical research 30, 1037–1043.

Ryberg, E., Larsson, N., Sjögren, S., Hjorth, S., Hermansson, N.O., Leonova, J., Elebring, T., Nilsson, K., Drmota, T., and Greasley, P.J. (2007). The orphan receptor GPR55 is a novel cannabinoid receptor. British journal of pharmacology 152, 1092–1101.

Scholz, H., Franz, M., and Heberlein, U. (2005). The hangover gene defines a stress pathway required for ethanol tolerance development. Nature 436, 845–847.

Scholz, H., Ramond, J., Singh, C.M., and Heberlein, U. (2000). Functional Ethanol Tolerance in Drosophila. Neuron 28, 261–271.

Singh, C.M., and Heberlein, U. (2000). Genetic control of acute ethanol-induced behaviors in Drosophila. Alcoholism, clinical and experimental research 24, 1127–1136.

Subbanna, S., Shivakumar, M., Psychoyos, D., Xie, S., and Basavarajappa, B.S. (2013). Anandamide–CB1 Receptor Signaling Contributes to Postnatal Ethanol-Induced Neonatal Neurodegeneration, Adult Synaptic, and Memory Deficits. The Journal of Neuroscience 33, 6350.

Turna, J., Syan, S.K., Frey, B.N., Rush, B., Costello, M.J., Weiss, M., and MacKillop, J. (2019). Cannabidiol as a Novel Candidate Alcohol Use Disorder Pharmacotherapy: A Systematic Review. Alcoholism: Clinical and Experimental Research 43, 550–563.

Vinod, K.Y., Yalamanchili, R., Xie, S., Cooper, T.B., and Hungund, B.L. (2006). Effect of chronic ethanol exposure and its withdrawal on the endocannabinoid system. Neurochemistry International 49, 619–625.

Viudez-Martínez, A., García-Gutiérrez, M.S., Navarrón, C.M., Morales-Calero, M.I., Navarrete, F., Torres-Suárez, A.I., and Manzanares, J. (2018). Cannabidiol reduces ethanol consumption, motivation and relapse in mice. Addiction biology 23, 154–164.

Wang, L., Liu, J., Harvey-White, J., Zimmer, A., and Kunos, G. (2003). Endocannabinoid signaling via cannabinoid receptor 1 is involved in ethanol preference and its age-dependent decline in mice. Proc Natl Acad Sci U S A 100, 1393–1398.

Wolf, F.W., Rodan, A.R., Tsai, L.T., and Heberlein, U. (2002). High-resolution analysis of ethanol-induced locomotor stimulation in Drosophila. The Journal of neuroscience : the official journal of the Society for Neuroscience 22, 11035–11044.

Young-Wolff, K.C., Tucker, L.-Y., Alexeeff, S., Armstrong, M.A., Conway, A., Weisner, C., and Goler, N. (2017). Trends in Self-reported and Biochemically Tested Marijuana Use Among Pregnant Females in California From 2009-2016. JAMA 318, 2490–2491.

